# Improvement of cryo-EM maps by density modification

**DOI:** 10.1101/845032

**Authors:** Thomas C. Terwilliger, Steven J. Ludtke, Randy J. Read, Paul D. Adams, Pavel V. Afonine

**Affiliations:** Los Alamos National Laboratory, Los Alamos NM 87545 USA; New Mexico Consortium, Los Alamos NM 87544 USA; Baylor College of Medicine, Houston, TX 77030 USA; Cambridge Institute for Medical Research, Cambridge, CB2 0XY, UK; Molecular Biophysics & Integrated Bioimaging Division, Lawrence Berkeley National Laboratory, Berkeley, CA 94720-8235, USA; Department of Bioengineering, University of California Berkeley, Berkeley, CA, USA

## Abstract

A density modification procedure for improving maps produced by single-particle electron cryo-microscopy is presented. The theoretical basis of the method is identical to that of maximum-likelihood density modification, previously used to improve maps from macromolecular X-ray crystallography. Two key differences from applications in crystallography are that the errors in Fourier coefficients are largely in the phases in crystallography but in both phases and amplitudes in electron cryo-microscopy, and that half-maps with independent errors are available in electron cryo-microscopy. These differences lead to a distinct approach for combination of information from starting maps with information obtained in the density modification process. The applicability of density modification theory to electron cryo-microscopy was evaluated using half-maps for apoferritin at a resolution of 3.1 Å and a matched 1.8 Å reference map. Error estimates for the map obtained by density modification were found to closely agree with true errors as estimated by comparison with the reference map. The density modification procedure was applied to a set of 104 datasets where half-maps, a full map and a model all had been deposited. The procedure improved map-model correlation and increased the visibility of details in the maps. The procedure requires two unmasked half-maps and a sequence file or other source of information on the volume of the macromolecule that has been imaged.

---

Single-particle electron cryo-microscopy (cryo-EM) is rapidly becoming the dominant technique for determination of large three-dimensional structures of macromolecules and their complexes^1^. The result of a cryo-EM analysis is a three-dimensional map reflecting the electric potential of the macromolecule^2^ and which has map values and an appearance closely related to maps obtained from X-ray crystallography^3^. In both cryo-EM and in macromolecular crystallography, the accuracy of the map is an important characteristic. In macromolecular crystallography, the amplitudes of Fourier coefficients are generally measured accurately and the phases are poorly estimated. It is common practice in that field to carry out a procedure known as density modification to reduce the errors in the phases and thereby improve the resulting map^4-7^. The source of new information in crystallographic density modification is prior knowledge about expected values in a map. For example, the probability distribution of map values may be known, or there may be knowledge about specific features in the map such as a flat solvent region. Information about the true density in part or all of the map can be used to obtain improved estimates of the phases, and these improved phases lead to an improved map everywhere, not just where the information was applied^6^.

In cryo-EM a form of density modification may be applied during the process of image reconstruction. The macromolecule typically occupies only a small part of the volume of the reconstruction, and during reconstruction noise is removed from the part of the map that is outside the macromolecule^8,9^. This can improve the map in the region of the macromolecule and is related to the “solvent flattening” aspect of crystallographic density modification^10^. Local denoising^11^ or filtering^12,13^ procedures are often applied to improve the interpretability of cryo-EM maps. A procedure for histogram-matching and resolution filtering has also been developed^14^. Though the overall process of density modification as implemented in a crystallographic setting is thought to be inappropriate for cryo-EM^15^, it has been suggested that the general concept could be adapted and applied^8,15^. Here we show that a version of density modification with the same theoretical basis as crystallographic density modification but with key differences reflecting the differences between crystallographic and cryo-EM maps can be used to improve cryo-EM maps.

There are several frameworks for density modification that could be applied to cryo-EM maps^4^. Here we use maximum-likelihood density modification, as it makes a clear distinction between information coming from the original data and information that comes from expectations about the features in the map^5^. The process for map improvement by maximum-likelihood density modification involves identifying how the current Fourier coefficients can be changed so as to increase the plausibility of the map (expressed as a likelihood), while retaining compatibility with the original experimental map (see Methods).

Cryo-EM maps differ in fundamental ways from crystallographic maps, and the actual process of density modification cannot be applied in the same way in the two situations. As detailed in Methods, one key difference is that in crystallography only the amplitudes of Fourier coefficients are measured directly (phases are generally estimated indirectly by comparison of amplitudes measured at different wavelengths or from slightly differing crystals), while in electron cryo-microscopy both phase and amplitudes are directly available from experiment. Another is that half-maps with errors that are nominally independent are available in electron cryo-microscopy^16^ but not in crystallography.

We tested the applicability of density modification theory to cryo-EM by applying it to a map of apoferritin at a reported resolution of 3.1 Å (EM Data Bank entry EMD-20028^17^). We used a matched 1.8 Å reference map (EMD-20026) to evaluate the error estimates that make up a key part of the density modification process and to test the effects of density modification on map quality.

The density modification procedure for the 3.1 Å map was to be carried out using Fourier coefficients to a resolution of 2.5 Å, so we first checked the accuracy of the 1.8 Å map up to this resolution by calculating the Fourier shell correlation (FSC) of independent half-maps^16,18^. Fig. 1A shows that the two half-maps corresponding to the full 1.8 Å map have an FSC value above 0.97 at all inverse resolutions of up to 0.4 Å^-1^ (corresponding to a resolution of 2.5 Å). Half-dataset FSC values can be used to estimate the expected correlation of Fourier coefficients for a map to Fourier coefficients representing the true map using the formula^18^,

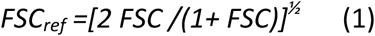

**Figure 1.**
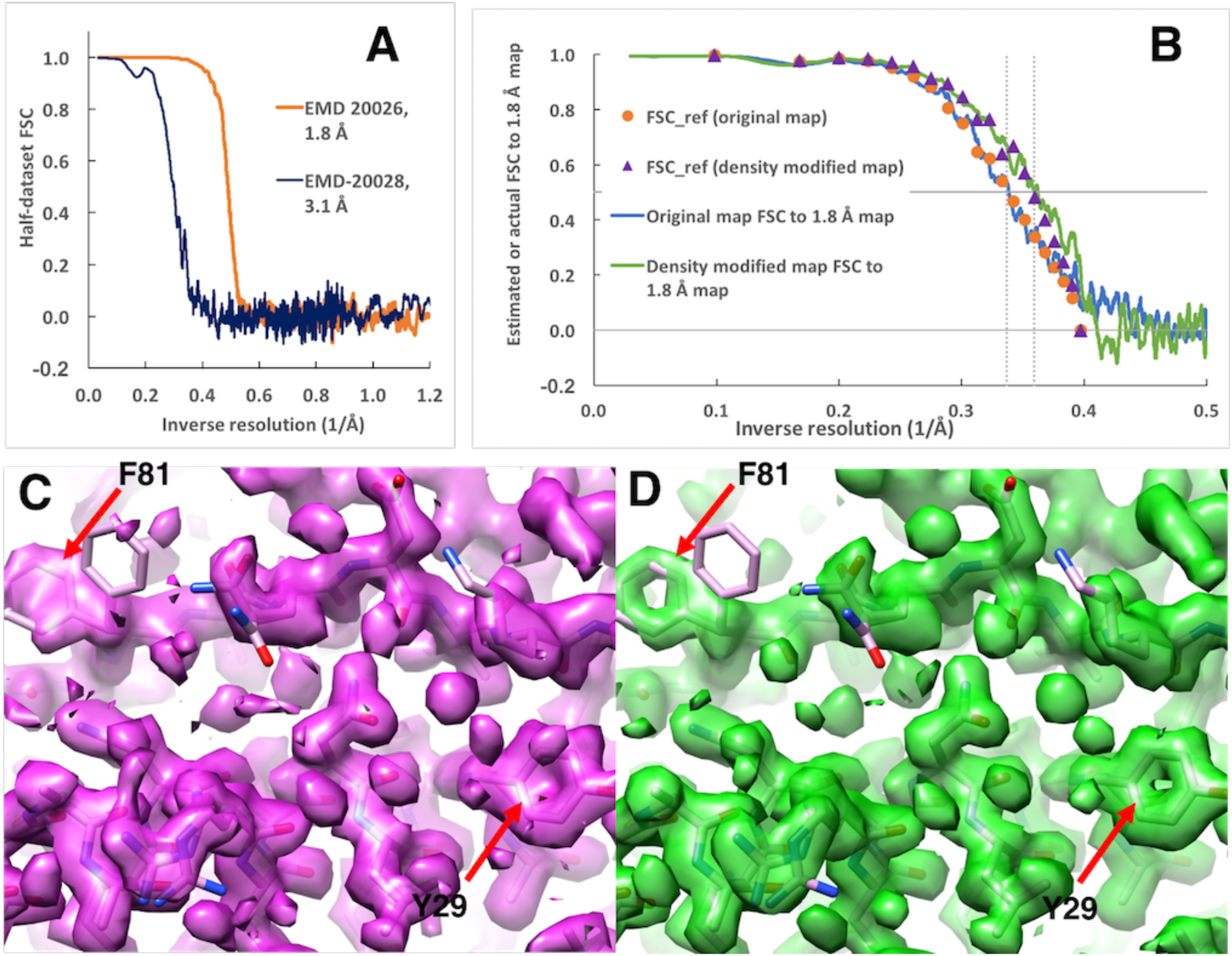
Density modification of apoferritin 3.1 Å map and evaluation using apoferritin 1.8 Å map. A. Fourier shell correlation (FSC) curves for EMD-20028 (3.1 Å) and EMD-20026 (1.8 Å). B. Orange circles are estimated resolution-dependent map quality (*FSC*_*ref*_) of original 3.1 Å map (see text). Blue line is the FSC of the original 3.1 Å map to the 1.8 Å map. Purple triangles are *FSC*_*ref*_ of density-modified 3.1 Å map, and the green line is the FSC of the density-modified 3.1 Å map to 1.8 Å map. Horizontal lines are drawn at FSC values of zero and ½. Vertical dotted lines are drawn at inverse resolutions of 1/2.78 Å^-1^ and 1/2.97 Å^-1^. C, D. Density modification of apoferritin 1.8 Å map, carrying out density modification without auto-sharpening the starting half-maps and using a density-modification resolution of 1.8 Å. C. Deposited map with resolution-dependence matched to that of the density-modified map using the *Phenix*^20^ tool *auto_sharpen* with the *external_map_sharpening* method. D. Density-modified map. The estimated improvement in resolution where *FSC*_*ref*_ is ½ is 0.05 Å. Contours in C and D are drawn to enclose equal volumes^21^. The X-ray structure of human apoferritin (PDB entry 3ajo^22^) is shown after docking in the deposited map and re-refinement against the density –modified map. Arrows indicate the locations of F81 (multiple conformations in 3ajo) and Y27 where the aromatic rings are clear after density modification.

Here we use the notation *FSC*_*ref*_ to emphasize that this is a FSC-based resolution-dependent estimate of map similarity to a reference (ideal) map. This usage corresponds to the previous use^18^ of *C*_*ref*_ and is similar to the use of the notation *CC\** in crystallography^19^). According to Eq. (1), an FSC value of 0.97 corresponds to a value of *FSC*_*ref*_ =0.99, indicating (aside from systematic errors affecting both half-maps^15^) that up to a resolution of 2.5 Å, the 1.8 Å map closely matches a perfect map of this structure.

We applied our density modification procedure to the two half-maps from the 3.1 Å dataset, yielding two intermediate map-phasing half-maps, two density-modified half-maps, and a final density-modified full map. As described in Methods, each Fourier coefficient in each map-phasing half-map is obtained individually by adjusting it to maximize the likelihood (believability) of a map calculated from this coefficient plus all other (constant) Fourier coefficients in the corresponding original half-map. These map-phasing half-maps are then recombined with the original half-maps using a resolution-dependent weighting approach to yield density-modified half-maps. Finally the density-modified half-maps are averaged to produce a single density-modified map. As a part of the density modification process, estimates are obtained of the Fourier shell correlations *FSC*_*ref*_ between the initial full map and a true map, and also estimated correlations *FSC*_*ref*_ between the density-modified map and a true map. The estimated Fourier shell correlation between the initial full map and a true map comes from applying Eq. (1) to a half-dataset FSC (such as the one shown in Fig. 1A for the 3.1 Å map). Fig. 1B illustrates these estimated resolution-dependent map accuracy values (*FSC*_*ref*_) for the 3.1 Å map (orange dots) and shows that they are very similar to actual map accuracy (the Fourier correlation between the 3.1 Å and reference 1.8 Å maps, blue line).

For the density-modified map, values of estimated Fourier shell correlation to a true map, *FSC*_*ref*_, are estimated from resolution-dependent error estimates (Eq. 9, Methods). These error estimates are used so that any correlations among the two original half maps and the two map-phasing half maps can be taken into account (see Methods). Fig. 1B displays estimates of resolution-dependent map quality *FSC*_*ref*_ for the density-modified map (purple triangles) and shows that they are very similar to the actual Fourier shell correlation values between the density-modified map and the 1.8 Å reference map (green line). Fig. 1B also shows that the values of both estimated Fourier correlation *FSC*_*ref*_ and actual Fourier correlation to the 1.8 Å reference map are consistently improved over the inverse resolution range from about 0.3 to 0.4 Å^-1^. This means that in the resolution range between about 2.5 Å and 3.3 Å the density modified map is more accurate than the original map.

In a real case, there will normally be no reference map for comparison. The analysis in Fig. 1B indicates that it is reasonable to use values of estimated Fourier correlation *FSC*_*ref*_, obtained from the correlations between original and density-modified half-datasets, as rough estimates of Fourier correlation to the true map.

Figs. 1C and D illustrate the visual effects of density modification of a high-resolution map, that of human apoferritin at 1.8 Å. The density-modified map in D shows high-resolution details that are not present in the resolution-matched map in C, including clear density for the aromatic rings of F81 and Y27. Taken as a whole, Fig. 1 indicates that application of density modification to apoferritin maps at 3.1 Å or 1.8 Å improves them in a significant way that is in general agreement with that expected from the error model for density modification described in Methods.

We next tested the generality of maximum-likelihood density modification of cryo-EM maps by applying it to 104 sets of half-maps and full maps available from the EM Databank (EMDB^17^). The main focus of this test was on maps in the resolution range of 2 Å to 4.5 Å where we developed the parameters and expected the procedure to work, but we included maps at lower resolution (up to 8 Å) as well. To evaluate the effects of density modification, we calculated the Fourier shell correlation^16^ of each map (before and after density modification) to the deposited atomic model from the Protein Data Bank (PDB^23^), after re-refining the model to the map to be evaluated (see Methods). The purpose of the re-refinement was to create similar biases for the analysis of original and density modified maps and yield a comparison that was fair. Using the map-model FSC values for an original and density-modified map, we generated two metrics that reflect different aspects of relative accuracies of the two maps. The first metric was the resolution at which the map-model FSC falls to approximately ½, an indication of a resolution where there is substantial information present, and the second was the average FSC (in the same resolution range for the two maps), a measure of overall quality of each map. Such metrics of map quality have many limitations^18,24-26^ and are therefore far less useful than a direct comparison with an essentially ideal map as in Fig. 1B. However, if they are fair they can at least give a general idea as to whether the method is useful.

Fig. 2A illustrates the resolution at which the map-model FSC falls to approximately ½ for original maps (blue dots) and density-modified maps (orange circles), plotted as a function of the reported resolution of the original maps. The resolution at which the map-model FSC falls to approximately ½ is generally improved (resulting in a smaller value) by density modification over the entire range of reported resolutions, though for maps with reported resolution worse than 4.5 Å the variability in this metric is quite large.

**Figure 2.**
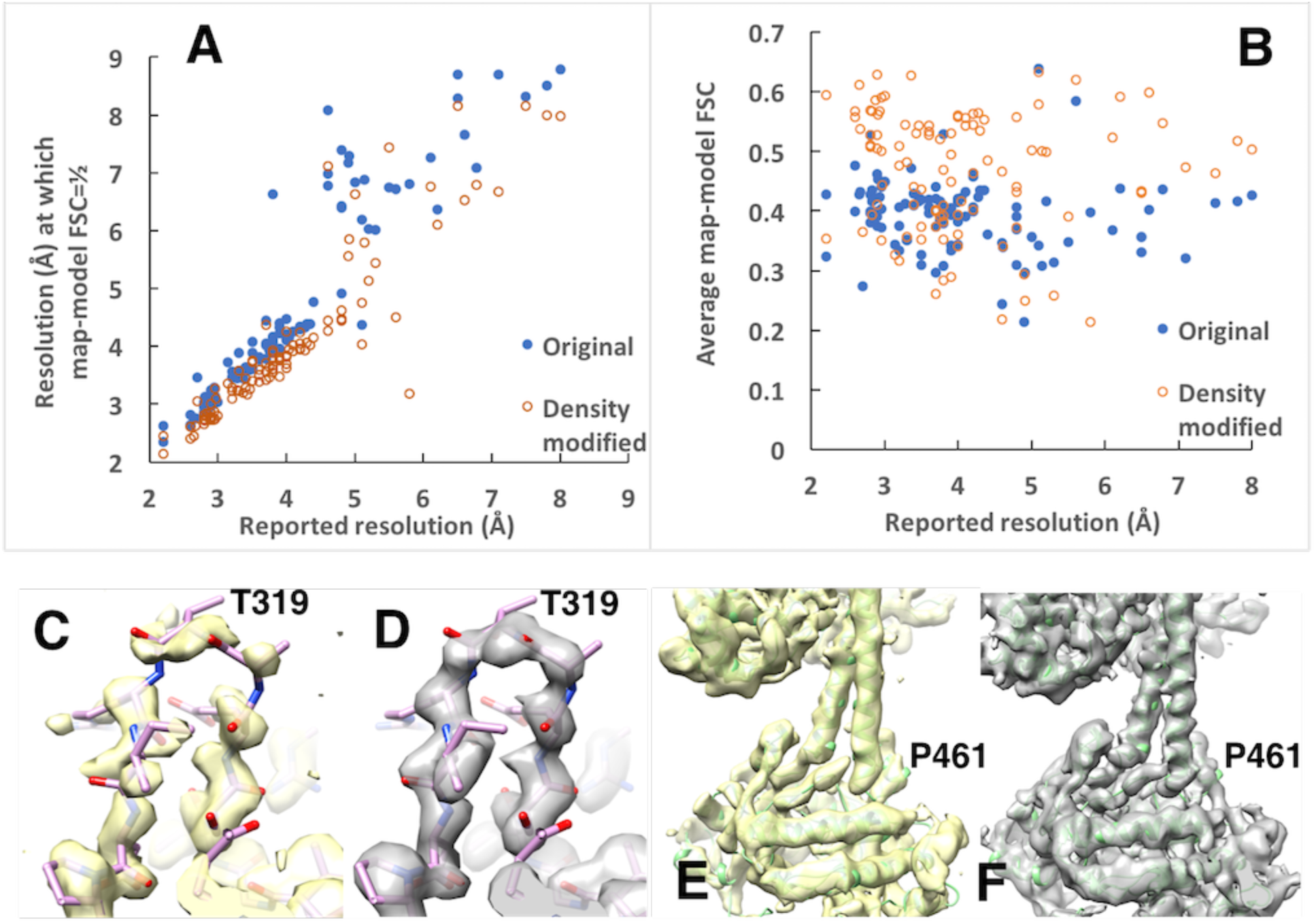
Application of density modification to maps from the EM Data Bank. A. Estimated resolution at which map-model FSC falls to approximately ½ for original maps (blue dots) and density-modified maps (orange dots) based on map-model FSC using models refined against the maps being examined. B. Mean map-model FSC for original maps (blue dots) and density-modified maps (orange dots), calculated up to inverse resolution of corresponding to 5/6 the stated resolution of the map (the value of 5/6 the map resolution is a typical value of the resolution used in density modification). C, D. Deposited and density-modified maps for β-galactosidase (2.2 Å, EMDB 2984). The resolution-dependence of the deposited map is matched to that of the density-modified map, and contours include equal volumes. Supplementary Fig. S1A shows the same region for the deposited map for EMDB 2984. E,F, Deposited and density-modified maps of a guanylate cyclase^28^ at 5.8 Å (EMDB entry 20282, PDB entry 6pas). Supplementary Fig. S1B shows the same region for a sharpened version of the deposited map.

Fig 2B shows the average map-model FSC for each original map (blue dots) and density-modified map (orange circles) as reflected in the mean map-model Fourier shell correlation, again plotted as a function of reported resolution. This metric also generally improves (is increased) after density modification but there is substantial variability in the amount of improvement (average improvement with density modification was 0.07, standard deviation of 0.08). Overall, both the resolution at which the map-model FSC falls to approximately ½ and average FSC generally improve with density modification but there is considerable variability, particularly at lower resolution.

To examine the visual effects of density modification in specific cases, Fig. 2 panels C-F show two matched pairs of deposited maps and corresponding density modified maps. To make the visual comparison as fair as possible, each deposited map was sharpened automatically to match the resolution-dependence of the density-modified map, and contours for matched maps were chosen to enclose equal volumes^21^ as in Fig 1. Figs. 2C and 2D show a loop region (residues A793-A804) that is poorly resolved in the β-galactosidase map at a resolution of 2.2 Å (EMDB entry 2984^27^; PDB entry 5a1a; Fig. 2C and Supplementary Fig. 1A), but is clear in the density-modified map (Fig. 2D). Figs. 2E and 2F show a 5.8 Å cryo-EM map of a guanylate cyclase^28^ (EMDB entry 20282, PDB entry 6pas). In the original map (Fig. 2E, resolution matched to the density-modified map) helices are essentially featureless tubes of density, while in the density-modified map (Fig. 2F) the helices have clear periodicity corresponding to the helical repeat. Sharpening of the original map does not yield a more interpretable map (Supplementary Fig. 1B). In each case the density modified map shows substantially more detail than the original map.

A limitation in our current implementation of density modification for cryo-EM is the assumption of relatively uniform noise levels throughout the region outside the macromolecule, while actual maps typically have noise levels that vary with distance from the macromolecule. As described in Methods, using a sub-volume (box) containing the macromolecule and a small region around it in density modification reduces the non-uniformity in noise levels but does not eliminate it. We investigated whether a reconstruction method that produced more uniform noise would improve density modification. We processed a subset of images available for β-galactosidase (EMPIAR-10061^27,29^) with two different procedures using EMAN2^30^, yielding maps with resolutions of about 3.9 Å. The first procedure was a standard reconstruction with default parameters including a Gaussian kernel with a resolution-dependent width except that no final masking was applied to the half-maps. A full map with masking was also generated from these half maps and used for comparison. The second procedure was a reconstruction with a fixed-width Gaussian kernel, expected to yield a more uniform noise distribution in real-space, to test whether the non-uniformity in noise in a cryo-EM map is a limiting factor. The maps obtained from each procedure were then density modified. Each map was then evaluated based on Fourier shell correlation to the deposited 2.2 Å map for β-galactosidase (EMD 2984^27^), superimposed on the reconstructions. The resolution at which this Fourier shell correlation falls to approximately ½ was used as a quality measure of the corresponding map.

Simple averaging of the half-maps from the standard reconstruction gave a map where the Fourier shell correlation falls to ½ at a resolution of 4.0 Å based on a comparison to the deposited 2.2 Å map. Density modification of these maps gave a map where it was 3.9 Å. Simple averaging of the half-maps with more uniform noise yielded a map for which the resolution where the Fourier shell correlation falls to ½ was 3.9 Å. This was improved by density modification to a value of 3.7 Å.

A portion of the initial full map is shown in Fig. 3A. For comparison, Fig. 3D shows the deposited high-resolution (2.2 Å) map, low-pass filtered at a resolution of 3.5 Å. The density-modified maps (Fig. 3B and 3C) both are more similar to the high-resolution, low-pass filtered map than the initial full map (Fig. 3A), with the map obtained by density modification of the half-maps with more uniform noise being the clearer of the two maps and the most similar to the high-resolution, low-pass filtered map (Fig 3C). The observation that the reconstruction protocol with relatively uniform noise produced the clearest map indicates that such a protocol may be particularly well-suited for density modification. In general, density modification is most likely to improve maps that have not been masked and in which the macromolecule is surrounded by a solvent region that retains the original noise from the reconstruction. The density modification process can then use that noise in the solvent to identify errors in Fourier coefficients and thereby to reduce noise in the region of the macromolecule.

**Figure 3.**
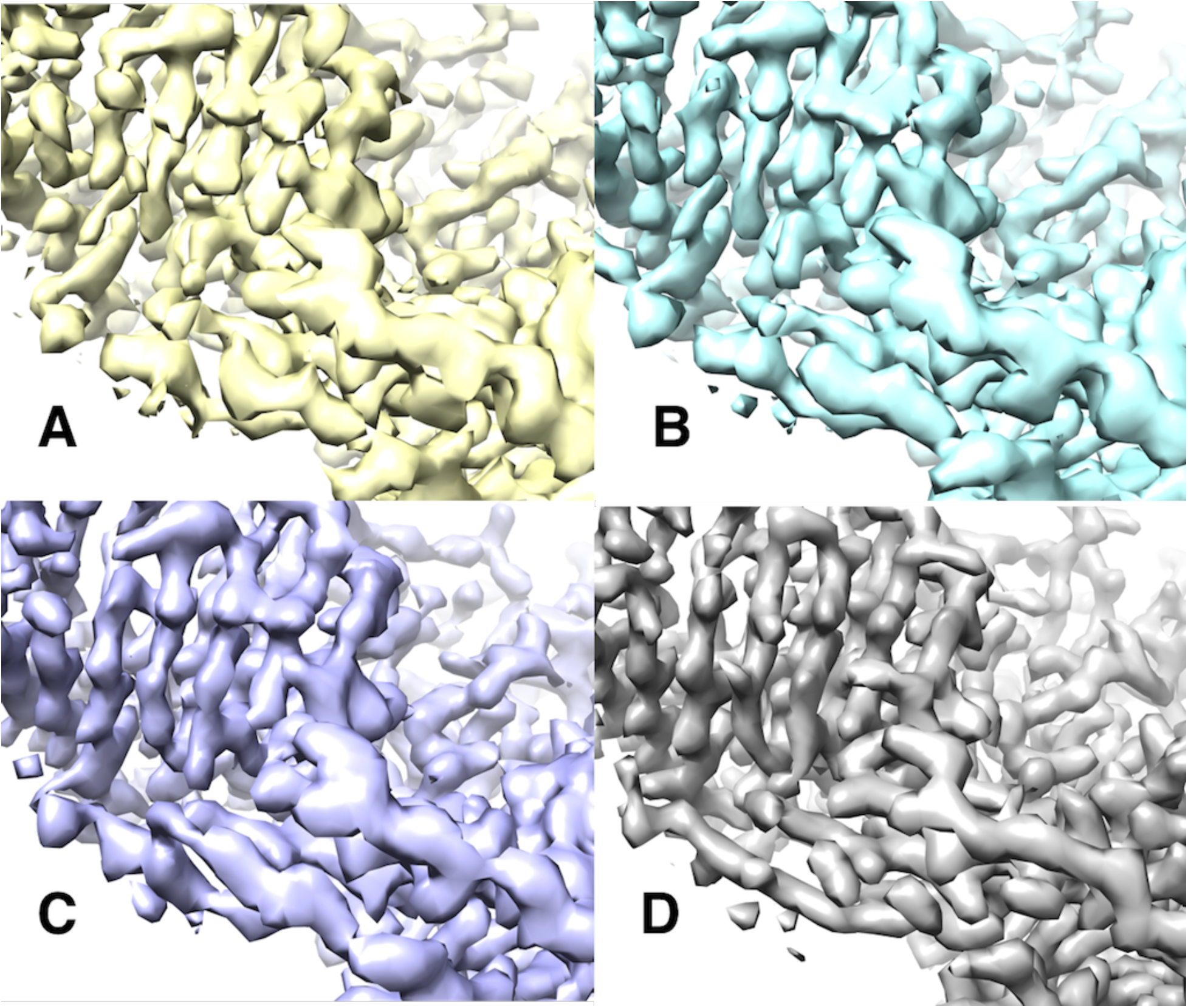
Effect of a reconstruction procedure yielding more uniform noise on the outcome of density modification. A. Standard reconstruction procedure applied to subset of images of for β-galactosidase (EMPIAR-10061). B. Density-modified version of the map in A. C. Density modified version of reconstruction designed to improve the uniformity of noise in the map. D. Deposited 2.2 Å map of β-galactosidase, superimposed on the map in A and low-pass filtered at a resolution of 3.5 Å. Each map was automatically sharpened using model-based sharpening with the deposited model for the 2.2 Å map of β-galactosidase (5a1a) superimposed on the map in A. Contours were chosen to enclose equal volumes in all four maps^21^.

The density modification procedure described here is fully automatic and requires only two half-maps, an optional nominal resolution, and information about the molecular volume of the macromolecule (such as a sequence file or molecular mass). Each of the 104 datasets analyzed in Fig. 2 took from 1 to 100 cpu-minutes (average of 12 minutes) on 2.3 GHz AMD processors. All of these analyses were carried out with default parameters in “quick” mode, supplying a sequence file and the reported resolution, and carrying out one cycle of density modification. In many of these cases, improved maps are obtained with additional cycles of density modification. In addition to a final density-modified map, the procedure yields two density-modified half-maps. As noted in Methods, these maps may have some correlations introduced by the masking effects of density modification, but with this caveat they can in principle be further processed with local sharpening^31^, weighted combination of half-maps and other methods for optimizing the final map.

There are numerous extensions to the methods that are described here that could improve the outcome of density modification. In particular, the analysis could include information about the macromolecule from other sources such as models built using the maps or fitted into them^32,33^. Density modification including symmetry not used in the reconstruction process could be carried out as well^34^. The procedure could allow starting half-maps that have errors that are correlated with each other or that have different expected errors, and errors that do not follow Gaussian distributions. It could be carried out using just one map or more than two ‘half-maps”. Maps could be density modified without boxing by introducing a location-dependent expectation for map values outside the macromolecule^4^. Errors could be estimated in regions of reciprocal space or anisotropically rather than in shells of resolution. The density modification step could also be carried out by other methods, for example solvent flipping^35^. The molecular composition could be calculated by analysis of the map, such as using local histograms of map values, allowing the identification of unexpected components. More generally, the entire analysis described here could be extended to any situation where a map of any dimensionality has errors that are at least partially independent in the Fourier domain and in which some information about expected values in the map is available. As suggested some time ago^8^, a density modification procedure such as the one described here could be incorporated into the overall process of iterative cryo-EM map refinement as well.

## Methods

### Maps and models

The maps used to generate Fig. 1 are apoferritin maps EMD 20026 and 20028 and their associated half-maps. These maps have reported resolutions of 1.75 Å and 3.08 Å, respectively and we refer to them as the “1.8 Å” and “3.1 Å” apoferritin maps. The estimates of resolution for these maps based on comparison of masked half-maps obtained in this work are slightly different, presumably due to different masking procedures, with values of 1.93 Å and 2.97 Å, respectively. The model shown in Fig 1 is derived from PDB entry 3ajo^22^ and has been superimposed on the 1.8 Å map and refined with the *Phenix* tool *real_space refine*^36^ against the density-modified 1.8 Å map.

The 104 sets of data used in Fig. 2 were chosen from the EMDB using data at resolutions from 2 Å – 8 Å that had associated half-datasets and matching models in the PDB.

Calculations of map-model Fourier shell correlations were done with a soft mask around the atoms in the model. The soft mask was calculated as a mask around each atom with a radius given by the atomic radius of the atom plus twice the resolution of the map, followed by smoothing of the map with a Gaussian smoothing function with a falloff to *1/e* of twice the resolution of the corresponding map. Analyses in Fig. 2 were carried out using the *Phenix*^20^ tool *resolve_cryo_em*, providing two half-maps, the reported resolution of the dataset, and the sequence of the macromolecule obtained from the deposited corresponding model. All 104 analyses were done using the same version (3689) and the same parameters except for resolution and sequence.

The data for Figures 1-3 are available as an Excel worksheet in Supplementary Data I, and the (sharpened) original and density-modified maps shown in Figs. 1-2 along with Chimera scripts to display them are available online at: http://phenix-online.org/phenix_data/terwilliger/denmod_2020/.

### Procedure for evaluation of map quality

We used an automated procedure to evaluate map accuracy and to choose matching map contours for display so that map comparisons would be as fair as possible. For evaluation of map accuracy we calculated Fourier shell correlations between a map and an atomic model refined against that map. The rationale for this procedure is that the atomic models available from the PDB are normally already refined against the deposited map. This necessarily biases the map-model FSC calculation. To make a comparison with a new map, the model is first re-refined against the original map. Then the re-refined model refined against the new map before FSC calculation. This is intended to lead to similar biases for models refined against original and density modified maps, leading to a relatively fair comparison between maps.

Our largely automated procedure for evaluation and display of one map was then (1) refinement of the corresponding model from the PDB using that map, (2) boxing the map with a rectangular box around the model with soft edges, (3) calculation of map-model FSC, (4) sharpening the map based on the map-model FSC^18,37^, (5) calculation of the resolution at which the map-model FSC falls to approximately ½, and (6) calculation of average map-model FSC^38^ up to a resolution of 5/6 the stated resolution of the map (i.e., *0.83 d*_*min*_). Then to compare a density-modified and original map visually, the maps were visually examined and a region of the map and contour level for the density-modified map were chosen where differences from the original map were clear. The contour level for the original map was then chosen automatically to yield the same enclosed volumes in the two maps^21^. This contour level for the original map was always close to that obtained by simply adjusting it to make about half the surface the color of the original map and half the color of the density-modified map when the two maps are displayed at the same time in Chimera^39^. Finally, keeping the same contour levels, the maps in Fig. 3 were masked 3 Å around the atoms in the region to be displayed to make it easier to see the region of interest. For the maps in Figs. 1 and 2, an additional step was added in which the resolution-dependence of the original map was matched to that of the density-modified map (using the *Phenix* tool *auto_sharpen* with the *external_map_sharpening* method) to make the comparison of maps as fair as possible.

### Errors in Fourier coefficients representing cryo-EM maps

We assume that the distribution of errors in Fourier coefficients representing cryo-EM maps can be represented by a two-dimensional Gaussian in the complex plane^8^. This assumption is evaluated in Fig. 4 which compares Fourier coefficients for apoferritin from the 3.1 Å and 1.8 Å maps analyzed in Fig. 1. Fourier coefficients for the shell of resolution from 3.0 Å to 3.1 Å were calculated for each map after boxing the maps around the fitted model used in Fig. 1. The Fourier coefficients for the 1.8 Å map were treated as perfect values. These values were multiplied by the correlation coefficient between the two sets of Fourier coefficients and subtracted from the Fourier coefficients from the 3.1 Å map to yield estimates of the errors in the 3.1 Å map. Fig. 4 shows histograms of these errors along directions parallel and perpendicular to the Fourier coefficients from the 3.1 Å map. In each case a Gaussian distribution is fitted to these histograms and is shown as well. It can be seen that the errors are not quite Gaussian and are not quite the same in the two directions, but that a Gaussian is a good first approximation. In this example, the normalized errors perpendicular to the Fourier coefficients from the 3.1 Å map have a mean of zero and a standard deviation of 0.63, while those parallel have a mean of 0.1 and a standard deviation of 0.70.

**Fig. 4.**
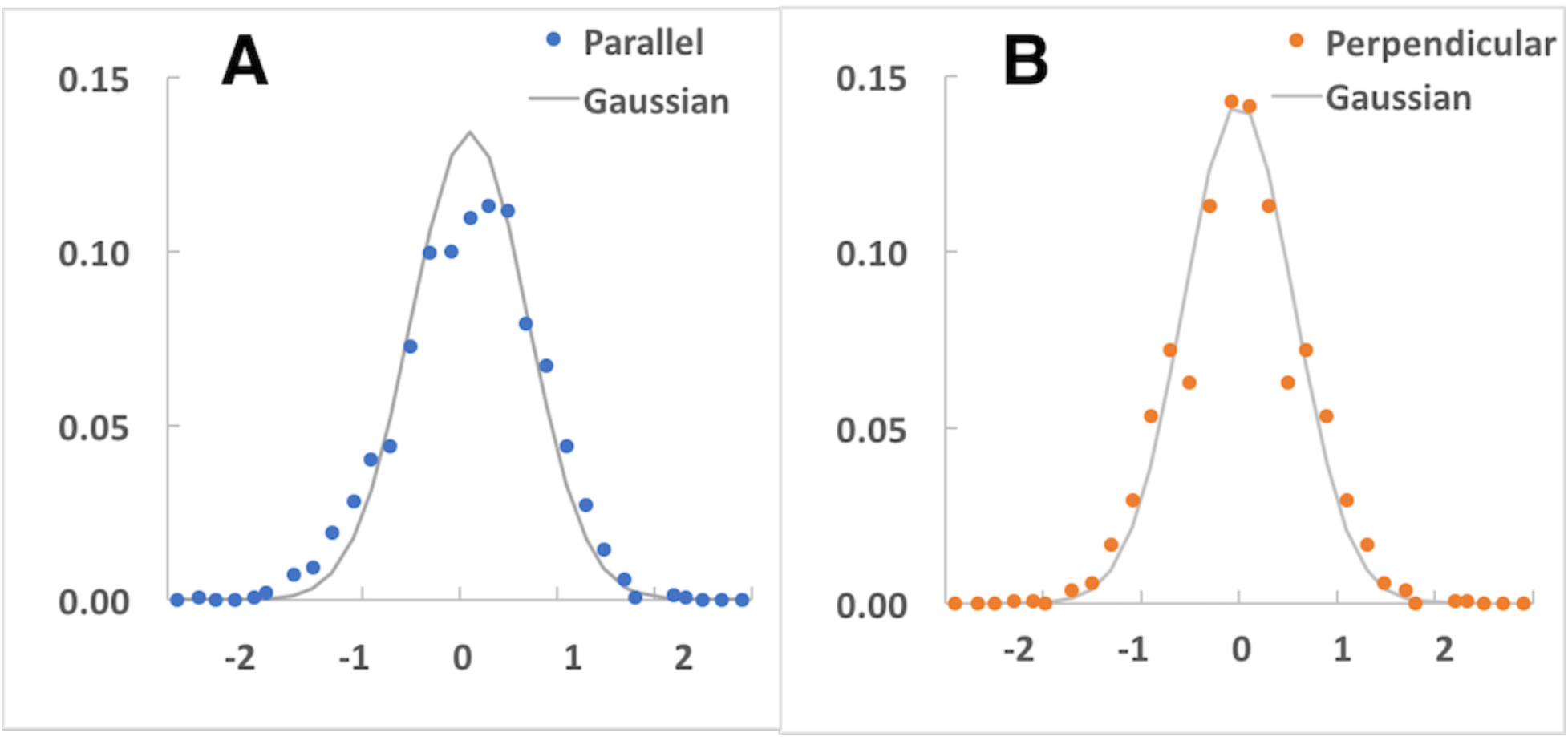
Analysis of distribution of errors in Fourier coefficients for apoferritin 3.1 Å map. A. Estimated errors in Fourier coefficients parallel to coefficients for 3.1 Å map. B. Errors perpendicular to coefficients for 3.1 Å map. Solid lines in each case correspond to a Gaussian fitted to the values shown (see text).

### Procedure for density modification of cryo-EM maps

Density modification of a cryo-EM map is based on the maximum-likelihood formalism that we developed previously for crystallographic density modification^5^. There are two important differences, however. One is that the starting probability distributions for Fourier coefficients (those available before density modification) are very different in the two cases. The other is that typically in a cryo-EM experiment, two independent half-maps are available (two maps with errors that are uncorrelated).

Maximum-likelihood density modification has two overall steps. In the first step a probability distribution (called the “map-phasing” probability distribution) is obtained for each Fourier coefficient. This map-phasing distribution has errors which, in an optimal situation, are independent of the errors in the corresponding Fourier coefficient in the starting map. In the second step the map-phasing probability distribution for each Fourier coefficient is recombined with the starting information about that Fourier coefficient to yield a new “density modified” estimate of that Fourier coefficient.

The first stage is the same in crystallographic and cryo-EM cases. It starts with a map represented by Fourier coefficients. It requires a function that describes how the likelihood (believability) of that map would change if the values in the map change^5^. This likelihood function might for example say that if the values in the map in the region outside the macromolecule all move towards a common value, the believability increases. It might also say that if the distribution of values in the region of the macromolecule becomes closer to an idealized distribution, that map’s believability improves. A specific example of a likelihood function that has both these properties has been described^5^ (Eq. 17 in this reference). Given such a map and likelihood function, it is possible to calculate a “map-phasing” probability distribution and its maximum or weighted mean for each Fourier coefficient^40^. This yields a “map-phasing” map.

The map-phasing map has the important property that the new estimate of a Fourier coefficient does not depend at all on the value of that Fourier coefficient in the starting map^40^. This rather non-intuitive situation is possible because the map-phasing probability distribution for a particular Fourier coefficient comes only from all the other Fourier coefficients and the characteristics of the map as reflected in the likelihood function. In other methods of density modification such as solvent flipping a similar effect is obtained by specifically removing the information corresponding to the original map^4,35^. It should be noted however that if the other Fourier coefficients have information about the errors in the Fourier coefficient in question (for example through previous density modification or by masking of the map around the molecule), that information can indeed affect the map-phasing estimate of the Fourier coefficient of interest.

The key differences in implementation between crystallographic and cryo-EM cases arise in the second step. First, the information about the Fourier coefficient that is available before density modification is very different in the two cases. In the crystallographic case, the amplitude of each Fourier coefficient is typically known quite accurately (often in the range of 5-30% uncertainty) and there may be some information about the phase (this might range from no information to phase uncertainties in the range of 45 degrees). The resulting distribution of likely values for a particular Fourier coefficient might essentially be a ring of relatively constant amplitude or a “boomerang” with partially-defined phase and relatively constant amplitude. In contrast, in the cryo-EM case, phase and amplitude are both uncertain, and the distribution of likely values before density modification can be represented by a two-dimensional Gaussian in the complex plane^8^.

This qualitative and very substantial difference in the form of the distribution of likely values for a Fourier coefficient prior to density modification means that when recombining information between the starting map and map-phasing distributions, different approaches are best suited to the two situations. For crystallographic applications, recombination essentially amounts to testing possible values of the phase of a Fourier coefficient at constant amplitude for consistency with prior and map-phasing distributions. In contrast, for cryo-EM applications as described here, recombination consists of calculating the product of two 2-dimensional distributions and finding the maximum of the resulting distribution. If the distributions are Gaussian, this amounts to a simple weighted average of the prior and map-phasing Fourier coefficients (cf. Eq. 7a below).

The second key difference is that cryo-EM analyses are typically carried out in a way that yields two half-maps with largely independent errors. This means that overall mean-square values of errors can be estimated in a straightforward way in bins or shells of resolution by comparison of Fourier coefficients from the two half-maps and from the two map-phasing half maps (see below).

The overall procedure for density modification of two half-maps is then (Supplementary Fig. S8): (1) average the two half maps and sharpen/blur the resulting map based on resolution-dependent half-map correlation^18,37^ to obtain an optimized starting full map, (2) identify regions in the full map corresponding to macromolecule or solvent (see below), (3) create or choose target histograms (see below) for the macromolecule and non-macromolecule regions, (4) use the histograms and Fourier coefficients representing each half-map in the first step of density modification to obtain a map-phasing probability distribution for each Fourier coefficient for that half-map, and (5) calculate a weighted average of values of each Fourier coefficient obtained from the two starting half-maps and the map-phasing maps obtained from them, yielding (6) a “density-modified” map along with corresponding weighted half-maps and resolution-dependent estimates of the accuracies and resolution of each map (Eq. 7a, below). The optimal weighting is discussed below in terms of a simple error model. Finally (7) the entire process can be repeated, using the density modified half maps from one cycle in step (4) of the next cycle. It is also possible (but not done by default in our current procedure) to use the full density modified map from one cycle to obtain histograms for the next cycle. The process is concluded after a specified number of cycles. For the examples in this work only one cycle was carried out (additional cycles did improve density modification for many of the 104 cases in Fig. 2, see Supplementary Fig. S7).

One variation on the overall procedure can sometimes improve density modification. This is to carry out density modification starting directly with the deposited maps instead of auto-sharpening them^37^ in step (1) above. This is controlled by the keyword *density_modify_unsharpened_maps=True* in the *Phenix* tool *resolve_cryo_em*.

### Identification of regions in a map corresponding to macromolecule and solvent

The regions in a map that correspond to macromolecule (as opposed to solvent) are identified as regions that have relatively high local variation in map values. The expected total volume occupied by the macromolecule is also considered, but not used as a fixed target as some parts of the macromolecule could be highly flexible (and therefore highly blurred) and as some other components (e.g., lipid) could also be present. The procedure is based on our method for probabilistic identification of the region occupied by a macromolecule in the unit cell of a crystal^41^. In essence, a density map is offset to set the mean value to zero, squared, and then smoothed with a smoothing radius of about twice the resolution of the map. The smoothed-squared map, Z(x) in our previous work^41^ then has high values where the density map has high variability, and low values where the density map is relatively flat. A Bayesian estimate of the probability that a particular grid point is in the region of the macromolecule is then calculated using the smoothed-squared map and using the approximate volumes occupied by macromolecule and solvent as priors^41^. Finally, a binary mask showing the macromolecule is obtained by choosing all points with probabilities of being inside the macromolecule higher than 50%. We note that this procedure has not been optimized. For example, setting the mean value to zero may not be the optimal approach. An improved boundary might instead be obtained in an iterative fashion by subtracting the mean value in the solvent region near the macromolecule^42^ from all points in the map. Additionally we did not test whether values other than the 50% cutoff would lead to improved solvent boundaries.

Note that the volume of the region identified as macromolecule in this procedure is not necessarily equal to the starting estimate of this volume. In practice, we find that a reasonable estimate of macromolecular volume can often be obtained simply by guessing several starting value of macromolecular volume, obtaining new estimates of macromolecular volume by applying the above procedure, and iterating the process to convergence.

### Target histograms of density distributions

A key element of the maximum-likelihood density modification procedure is the use of target histograms representing the expected distribution of map values for the “true” (desired) map in the region containing the macromolecule and in the region outside it^5,14^. These histograms can be obtained in any of several ways. One is to use a map or maps corresponding to high-quality structures that are already determined. Another is to use histograms used previously for crystallographic analyses. The default method used here is to use histograms based on crystal structures, with an option to use histograms derived from the full map obtained by averaging the two current half-maps. These averaged histograms have the advantage that they are automatically at the correct resolution and represent macromolecule and surrounding region in just the same way as the half map to be analyzed, but in our tests the histograms from crystallographic analyses resulted in the largest improvements so these are used by default.

Error model for analysis of FSC curves and its use in optimizing weights and estimating correlations to true maps.

We use a simple error model with the following assumption:

A. Starting half-maps have errors that are uncorrelated between half-maps. This assumption is based on the construction of half-maps, in which they are derived from independent subsets of the data. There are however some aspects of half-map construction that could lead to correlation of errors, including the use of the same reference in some stages of analysis and masking of the maps^18^.
B. Map-phasing half-maps have errors that are uncorrelated between half-maps and they may also have errors that are correlated with the corresponding starting half-maps and errors that are correlated with the each other. Errors correlated with the corresponding starting half-maps could come from the density modification procedure not yielding fully independent information. Errors correlated with the other density-modified half-map could come from masking effects introduced from solvent flattening procedures in density modification.
C. Mean square values of errors are resolution-dependent. This assumption simplifies the analysis of errors by describing the errors in terms of resolution and allowing them to be estimated in shells of resolution.
D. Mean square errors for each member of a pair of half maps are the same. This assumption comes from the construction of half-maps, where they typically come from equal numbers of images.
E. Errors have two-dimensional Gaussian distributions with mean expected values of zero. This simplifies the analysis.
F. Fourier coefficients representing individual half-maps are equal to the true Fourier coefficients plus uncorrelated errors unique to that map and correlated errors shared among two or more maps. This yields a simple form for the Fourier coefficients that is amenable to estimation of errors from correlations of Fourier coefficients.

These assumptions yield a simple error model where Fourier coefficients for the original and map-phasing half-maps can be represented as:

Original half-maps:

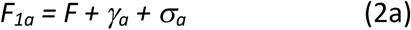

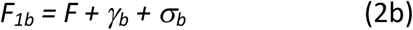

Map-phasing half-maps:

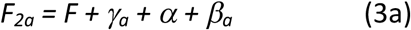

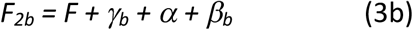

In this description, *F* represents the true value of one Fourier coefficient. There are estimates of F that come from each half-map and each map-phasing half-map. *F*_*1a*_ and *F*_*1b*_ represent Fourier coefficients for half-maps *a* and *b*, and *F*_*2a*_ and *F*_*2b*_ represent Fourier coefficients for map-phasing half-maps *a* and *b*. The terms *σ*_*a*_ and *σ*_*b*_ represent uncorrelated errors in half maps *F*_*1a*_ and *F*_*1b*_. The mean square values of each are *S*: *<σ*_*a*_^*2*^*> = <σ*_*b*_^*2*^*> = S*, and the mean values of all errors in this analysis are zero. The term *γ*_*a*_ represents errors that are correlated between half map *F*_*1a*_ and its corresponding map-phasing half map *F*_*2a*_ (present in half map *F*_*1a*_ and not corrected by map phasing), and the term *γ*_*b*_ represents errors correlated between *F*_*1b*_ and *F*_*2b*_. The mean square values of *γ*_*a*_ and *γ*_*b*_ are *<γ*_*a*_^*2*^*> = <γ*_*b*_^*2*^*> = C*. The terms *β*_*a*_ and *β*_*b*_ represent uncorrelated errors in half maps *F*_*2a*_ and *F*_*2b*_, where *<β*_*a*_^*2*^*> = <β*_*b*_^*2*^*> = B*. The term *α* represents errors correlated between half maps *F*_*2a*_ and *F*_*2b*_, where *<α*^*2*^*> = A*. As it is assumed that errors are resolution-dependent, the estimates of mean square errors (*A, B, C, S*) are in turn assumed to be resolution-dependent and in our procedure they are estimated in shells of resolution.

For simplicity in notation, we assume below that the Fourier coefficients for each half-map are normalized in such a way that the mean square value of F is unity. As the following calculations only involve correlation coefficients, the overall scale on Fourier coefficients has no effect on the values obtained, so this simplification does not affect the outcome of the analysis.

Using this error model and normalization, the expected values of correlations between half-maps can be calculated. These are as follows, where the brackets represent expected values, and the notation *CC(x,y)* represents the correlation coefficient relating values of *x* and *y*.

The expected correlation between half-maps *a* and *b* is given by,

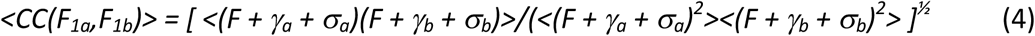

which reduces to,

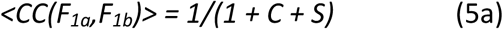

Similarly, correlation between map-phasing half-maps *a* and *b* is given by:

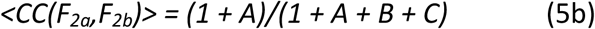

Cross-correlation between half map *a* and map-phasing half map *a* (and also between corresponding maps *b*) is given by:

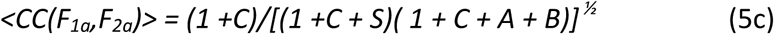

Cross-correlation between half map *a* and map-phasing half map *b* (and also between half map *b* and map-phasing half map *a*) is given by:

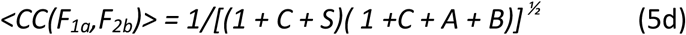

As there are four relationships and four parameters describing errors, the relationships can be used to estimate the values of the errors *A, B, C*, and *S*, leading to the formulas:

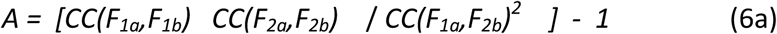

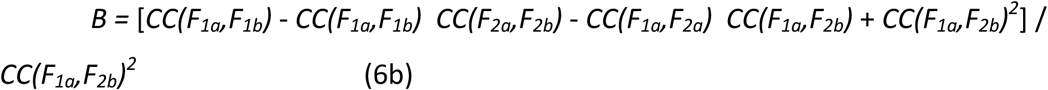

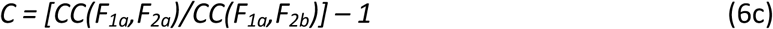

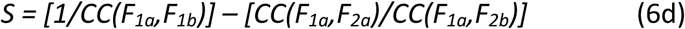

As noted in more detail below, for shells at high resolution the uncertainties in the correlations such as *CC(F*_*1a*_,*F*_*1b*_) can be large compared to the correlations themselves. In these situations the values of correlations are smoothed and additional assumptions are made about the relationships among the error estimates in order reduce the number of parameters that need to be obtained from the data at that resolution.

After estimation of errors, all estimates of *F* can be averaged with resolution-dependent weighting factors w. Based on the assumption of equal mean square errors in members of a pair of half-maps, the weights on each half map in a pair are always equal. The final density-modified Fourier coefficient (*G*) is obtained as the weighted average of the original and map-phasing map coefficients (*F*_*1*_ and *F*_*2*_) and is given by,

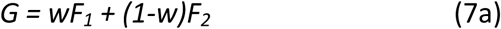

Where *w* is the weight on the original averaged half maps (*F*_*1*_) and *(1-w)* is the weight on averaged map-phasing half-maps (*F*_*2*_), and the averaged maps are given by:

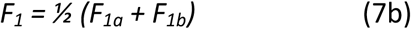

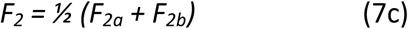

The weight *w* that maximizes the expected correlation of the estimated Fourier coefficient, *G* with the true one, *F*, is:

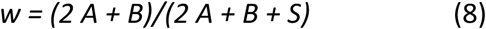

and the estimated correlation of *G* with *F*, the correlation of the final estimate of the Fourier coefficient with the true one, represented^18^ by *FSC*_*ref*_ is:

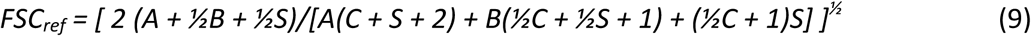

As mentioned above, assumptions are made to allow estimation of correlated and uncorrelated errors from FSC plots for resolution shells where uncertainties in correlation estimates are large. These additional assumptions are:

A. In the highest resolution shell considered there is no remaining signal and all correlations of Fourier coefficients are due to correlated errors. The limiting resolutions of FSC plots in this analysis are set in such a way that there is little signal at those resolutions and correlations are largely due to noise and correlated errors. FSC plots are calculated in shells of resolution (*d*). The highest resolution in error analyses considered (*d*_*min*_) is the resolution used for density modification multiplied by a fixed ratio (typically 5/6). As the resolution used for density modification is normally about 0.5-1 Å finer than the nominal resolution of the overall map, for a 4 Å map this high-resolution *d*_*min*_ would typically be in the range of 2.5 Å to 3 Å.
B. In high-resolution shells where there is substantial uncertainty in the estimates of errors (typically where half-map correlations are less than about 0.05), ratios of correlated to uncorrelated errors are assumed to be the same as those estimated in lower-resolution shells.
C. For shells of resolution (*d*) where the values of FSC are below a fixed minimum FSC, (typically *FSC*_*_min*_=0.2), smoothed FSC values are calculated by fitting the observed values to a simple exponential function with one free parameter. The function used is *FSC=(FSC*_*d1*_ - *FSC*_*d_min*_)*exp(-H/d*^*2*^*)+ FSC*_*d_min*_. The free parameter is *H*, the fall-off with *1/d*^*2*^). *FSC*_*d1*_ is the value of FSC at resolution *d1*, the highest resolution where FSC is higher than a fixed minimum value (typically 0.2). *FSC*_*d_min*_ is the estimated FSC at the highest resolution in the analysis. As noted above it is assumed that any non-zero FSC found at this resolution is due to correlated errors in the analysis.

### Real-space weighting and weighting of individual Fourier coefficients in calculation of the final map

An option available at the end of a cycle of density modification is to apply a local weighting scheme to the final combination of original and density-modified maps. The idea is to identify local accuracy in the original map from local similarity between original half-maps, and also local accuracy in the map-phasing maps from local similarity between map-phasing half maps. The procedure for error estimation for one pair of half-maps (of any kind) is to subtract the maps, square the resulting map, and smooth the squared map. The smoothing radius is typically given by twice the resolution of the map. This yields a local variance map with values approximately representing the sum of the variances of the two half-maps.

A local weight for each set of half-maps (original half-maps, *F*_*1a*_, *F*_*1b*_ and map-phasing half-maps, *F*_*2a*_, *F*_*2b*_) is then obtained as the inverse of the corresponding local variance map. These local weights are then scaled to yield an average local weight of unity and then are applied to the individual half-maps before they are averaged.

A second option for recombination of maps is to weight individual Fourier coefficients based on the estimated variance for these coefficients. The variance for an individual coefficient is estimated from the four Fourier coefficients representing the four half-maps available at the end of the procedure (the two original half-maps and the two density-modified half-maps). These two procedures are not applied by default but can improve maps in some cases.

### Overall spectral scaling

The procedure at this point yields weights (*w, 1-w*) for the two pairs of half-maps that are combined to yield a density-modified map, and an estimate of the quality of the map coefficients in this resolution shell, (*FSC*_*ref*_). This calculation is carried out in shells of resolution (typically 100 shells) and allows calculation of Fourier coefficients in each shell of resolution. A final resolution-dependent weighting (spectral scaling) is then applied to the Fourier coefficients. There are several options for this final scaling.

One option (the default, controlled by the keyword *final_scale_with_fsc_ref=True*) begins by applying a scale factor (*A*_j_) for each shell *j* to yield a constant value of average amplitudes of Fourier coefficients in each shell (essentially creating an E-map in crystallographic terminology; note that although scale factors are calculated in shells of resolution, they are applied as a smooth function of inverse resolution). We note that this procedure has a significant disadvantage in that the Fourier coefficients in the lowest-resolution shells typically have very high variation in magnitude compared to those in higher-resolution shells, so that no single scale factor really can be suitable. After this normalization step, Fourier coefficients are multiplied by the value of *FSC*_*ref*_ in that shell, corresponding to the approach often used to sharpen a map based on its corresponding half-maps^18^.

A second option for final scaling is similar to the first, except that in the first step the scale factor applied is the square root of the scale factor (*A*_j_) described above. This option is an attempt to reduce the effect of scaling on the low-resolution information and can improve the low-resolution map-model correlation found at the conclusion of density modification in some cases. This procedure is controlled by the keyword *geometric_scale=True* and is the default if *final_scale_with_fsc_ref=False.*

A third option, one that can be carried out after either of the first two, is to apply a resolution-dependent scale factor to all Fourier coefficients that corresponds to the resolution-dependence of a calculated “typical” protein structure (we used the 2.2 Å map EMD-2984 of β-galactosidase to calculate this resolution dependence). This is controlled by the keyword, *spectral_scaling=True* and is not carried out by default.

### Boxing of cryo-EM maps

In our procedure, a rectangular solid portion of a cryo-EM map that contains the macromolecule of interest is cut out from the map and is used in the analysis. This “boxing” of the map is carried out with a “soft” (Gaussian) mask with a smoothing radius typically equal to the resolution of the map to reduce the introduction of correlations in Fourier coefficients between different maps boxed in the same way^18^. The edges of the box are chosen using bounds in each direction identified using a low-resolution (typically 20 Å) mask calculated from the full map with a volume based on the expected molecular volume of the macromolecule. Typically, a buffer of 5 Å is added to the bounds in each direction to yield a box that has dimensions 10 Å bigger than the size in each direction of the macromolecule.

There are two important effects of boxing. One is to reduce the variation of noise in the map in the region outside the macromolecule. In a typical cryo-EM map there is substantial noise (fluctuation in map values not representing the macromolecule) near the macromolecule, and progressively less further from the macromolecule (the variation in noise levels may also be more complicated). This variation in noise levels is largely due to the use of procedures that smooth Fourier coefficients in reciprocal space with the effect of masking around the macromolecule^30^. In our procedure it is assumed that the distribution of map values in the region outside the macromolecule can be represented by a simple histogram. As there is a distance-dependent variation in the level of noise in unboxed cryo-EM maps, our procedure can be made more applicable by boxing the maps.

The second effect of boxing the map is to reduce the correlation of Fourier coefficients in the map. If a small object is placed in a large box and Fourier coefficients are calculated representing the object in the box, coefficients with similar indices (neighboring Fourier coefficients) will be highly correlated^43,44^. The significance of correlations between Fourier coefficients is that errors may be correlated as well, resulting in map-phasing Fourier coefficients that are not fully independent from the original Fourier coefficients. Boxing reduces the empty volume of the map and reduces this correlation.

### Effect of masking in density modification due to solvent flattening

An indirect effect of density modification is masking of the map. Density modification includes a step in which noise in the solvent region of the map is reduced, and this process can lead to a map that is partially masked. A consequence of full masking (setting values to a constant value outside the mask) is that correlations with a perfect map, with a model-based map, or between half-maps^16^ are increased relative to an unmasked map. We carried out a test to evaluate whether this effect contributes in a substantial way to the improvement in resolution by density modification shown in Fig. 1B. The test consisted of repeating the analysis in Fig. 1B except that instead of actually carrying out map phasing, each initial half-map was simply multiplied by the probabilistic mask that was to be used in identifying the protein region for density modification, and only one cycle was carried out (this can be done using the keyword *control_no_denmod* in the *Phenix* tool *resolve_cryo_em*). In Fig. 1B the resolution at which the map-model FSC falls to approximately ½ changed with density modification from 2.98 Å to 2.77 Å. In the test analysis using just masking, this value changed by only 0.01 Å (from 2.98 Å to 2.97 Å), indicating that the masking effect is very small in this case.

### Resolution cutoff used for density modification

In order to include information at high resolution, the resolution (*d*_*dm*_) of Fourier coefficients used in the density modification procedure is typically finer than the resolution of the full map. The relationship between the resolution *d* of a map and the optimal resolution *d*_*dm*_ for density modification is not obvious, so we used an analysis of 51 half-maps from the EMDB and associated models from the PDB to develop an empirical relationship. The empirical function was obtained by carrying out the entire density modification procedure for each dataset, each with a range of values of *d*_*dm*_. Then the average FSC between map and model-based map was calculated for each analysis and a simple function was developed for choosing the resolutions where density modification was optimal. This function, developed for resolutions between 2.4 Å and 5 Å, was:

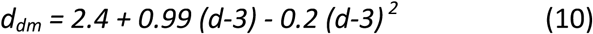

At a resolution *d=2.4* Å, this yields *d*_*dm*_ *= 1.9*. For maps with resolution finer than 2.4 Å, we simply subtract 0.5 Å from the resolution to yield *d*_*dm*_, except that *d*_*dm*_ is never allowed to be less than *½d*. For resolutions greater than 5 Å the optimal resolution for density modification has not been identified. In Fig. 2 Eq. (10) is used.

Three options for choice of resolution cutoff for density modification *d*_*dm*_ are available in the current implementation of density modification. One is directly specifying *d*_*dm*_, one is using Eq. 10 to estimate *d*_*dm*_, and the last is to try various values of *d*_*dm*_ and choose the one that leads to the most favorable estimated improvement in the resolution where *FSC*_*ref*_ is *½* based on Eq. 9.

### Adjustable parameters

There are many adjustable parameters in our procedure but all have default values (default values for version 3689 of *Phenix* were used in all 104 tests in Fig.2). Some of the parameters that can substantially affect the results and that a user might vary if the initial results are not optimal are listed here. The resolution used for density modification is not fully optimized and can affect the outcome substantially. Additional cycles of density modification can improve the outcome in many cases. The choice of final scaling procedure (*final_scale_with_fsc_ref*) can affect the density-modified map, as can the choice of half-map sharpening at the start of density modification (*density_modify_unsharpened_maps*). The number of shells of resolution used in the calculation of correlations between Fourier coefficients has a default of 100; more shells can potentially improve the accuracy by not grouping coefficients that have very different values simply due to resolution-dependent variation but could reduce it due to fewer coefficients in a calculation. The optional use of spectral scaling, real-space weighting, and individual weighting of Fourier coefficients at the end of the procedure can sometimes affect the resulting map.

### Histogram-matching

We examined whether map improvement can also be obtained by using histogram-matching^45^ without density modification. With histogram-matching alone, a new map value at each grid point is obtained based on the distributions of expected and observed values in the map and the value at that grid point. With histogram-matching as a part of density modification, this histogram-matched map is not used directly as the new map, but rather it is used as a likelihood target that indicates the plausibility of a candidate map. This allows information from all parts of the map to be combined to yield new estimates of each Fourier coefficient.

To test whether the map improvement obtained with density modification could be obtained using real-space methods alone, we applied histogram-matching^45^ to the 1.8 Å apoferritin map shown in Fig. 1C. Supplementary Fig. S2 illustrates that our histogram-matching approach improves this map, but density modification improves this map considerably more. Panel A shows the average of deposited half-maps, sharpened to match density-modified map in panel C. Panel B shows the histogram-matched version of the map in A, and panel C shows the density-modified map. Note the considerable improvement in clarity in the ring of F81 in the density-modified map.

### Effects of density modification on individual half-maps

Supplementary Fig. S3 shows the individual original half-maps and the two density-modified half maps produced by the 1.8 Å apoferritin density modification example shown in Supplementary Fig. S2. It can be seen that the original half maps have similar clarity but are different, and that both half maps are improved by density modification and are again somewhat different.

### Sensitivity of density modification to parameters

We tested the sensitivity of density modification to two parameters that seemed likely to have substantial effects on the procedure, the box size, and the resolution used for density modification. In each case we used the 3.1 Å apoferritin map analyzed in Fig. 1. Supplementary Fig. S4 shows the effect of varying the resolution used for density modification on the quality of density modification, as measured by the estimated improvement in resolution and actual improvement (change in the resolution where the map-model FSC was approximately ½). It can be seen that the density modification resolution has a small effect on the actual improvement over the range of 2 Å – 2.6 Å but, as expected, the improvement becomes smaller as the density modification resolution approaches the nominal resolution of the map. The estimated resolution has more variability but a similar relationship.

We investigated the effect of box size by carrying out density modification using different size boxes to extract the molecule, ranging from a box just the size of the molecule (defined as the region with expected volume where density is highest), to a box with edges 30 Å bigger in each direction (the default is a box 10 Å bigger than the molecule). Over this range of box sizes the resulting resolution where the FSC to the model equaled ½ varied by a small amount (ranging from 2.76 Å to 2.83 Å for all boxes except the smallest one which had a value of 2.92 Å), and with the best resolution of 2.76 Å with a box 4 Å bigger than the smallest one. This indicates that at least in this case the exact box size is not very important but that a small box may be slightly better than a bigger one.

### Changes in map-model metrics after density modification

We examined whether common model and map-model metrics were substantially different using density-modified maps compared to the original maps. Supplementary Fig. 5 shows that map-model correlation (calculated at a resolution given by 0.83 times the nominal resolution of the starting map to emphasize high-resolution information) generally improves, but that the other metrics examined (rotamer outliers, ClashScore^46^, Ramachandran % favored, EMRinger^47^ scores) did not change in a systematic way over the resolution range of 2 Å – 4.5 Å where our method is designed to apply.

Supplementary Fig. S6 shows the local map-model correlation for EMD-7544 and associated PDB entry 6coy^48^ before and after density modification. The density modified map has generally higher map-model correlation but the difference varies somewhat by location along the chain.

### Effect of applying multiple cycles of density modification

Supplementary Fig. S7 illustrates the effect of multiple cycles of density modification on the average map-model FSC obtained. Panel A corresponds to Fig. 2B after 5 cycles of density modification. Panel B compares the average map-model FSC after one cycle with the same metric after 5 cycles. It can be seen that additional cycles lead to a substantial improvement in a number of cases, while in many other cases there is little effect after additional cycles, and in 2 cases there is substantial worsening.

## Software availability

All the procedures described in this work are available using the *Phenix* tool *resolve_cryo_em* in versions 3689 and later of the *Phenix* software suite^20^.

## Acknowledgements

This work was supported by the NIH (grant GM063210 to PDA, RJR and TT and grant R01-GM080139 to SJL), the Wellcome Trust (grant 20947/Z/17/Z to RJR), and the Phenix Industrial Consortium. This work was supported in part by the US Department of Energy under Contract No. DE-AC02-05CH11231 at Lawrence Berkeley National Laboratory.

## Author contributions

SL carried out image processing of test datasets to evaluate varying reconstruction procedures, RJR and TCT contributed ideas on the form of errors in cryo-EM, PA developed tools for the testing infrastructure, TCT developed the software for error analysis, and PDA and TCT supervised the work.

## Author information

The authors declare no competing financial interests.

## Supplementary Figures

**Supplementary Fig. S1.**
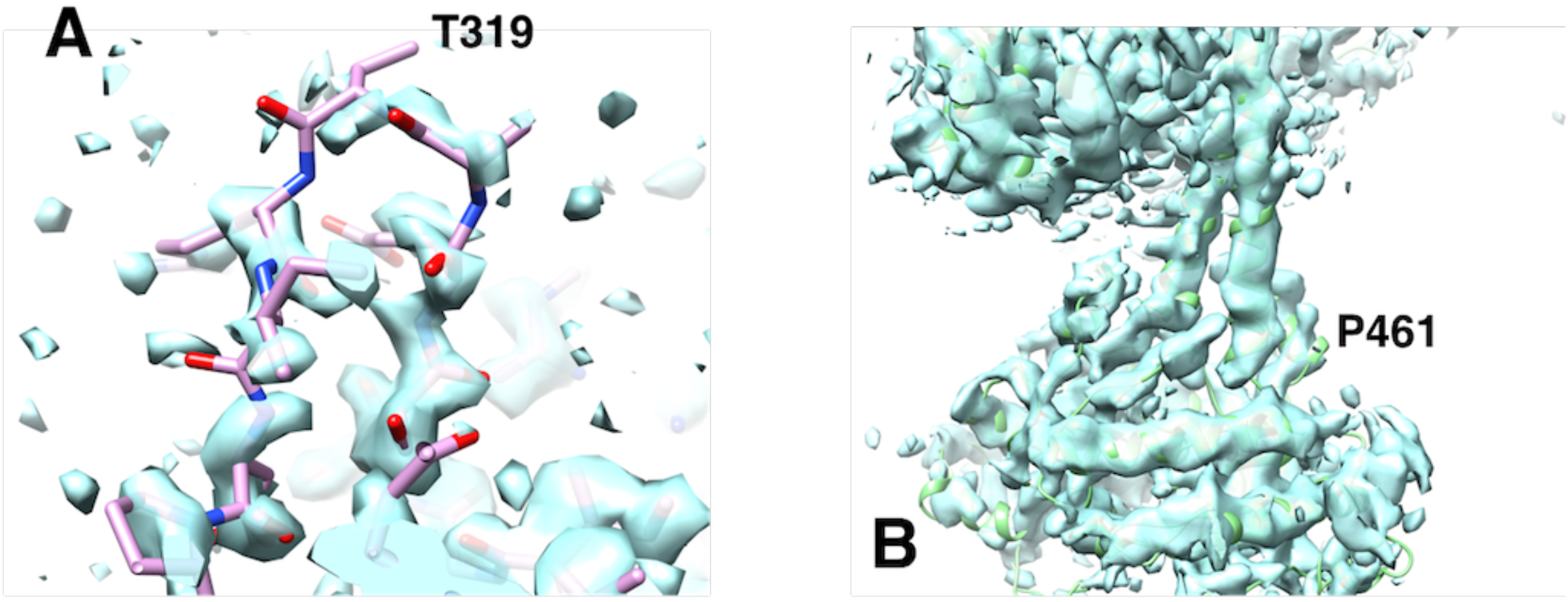
Analysis of non-density-modified maps to examine whether simple modifications (sharpening, using deposited maps) would yield maps that appear similar to the density-modified maps. A. Deposited map showing the same region as depicted in Fig. 2C. This map also shows poor density for the loop. Varying the sharpening of the map did not yield connected density as in Fig. 2D. B. Sharpened version of the original map shown in Fig. 2E.

**Supplementary Fig. S2.**
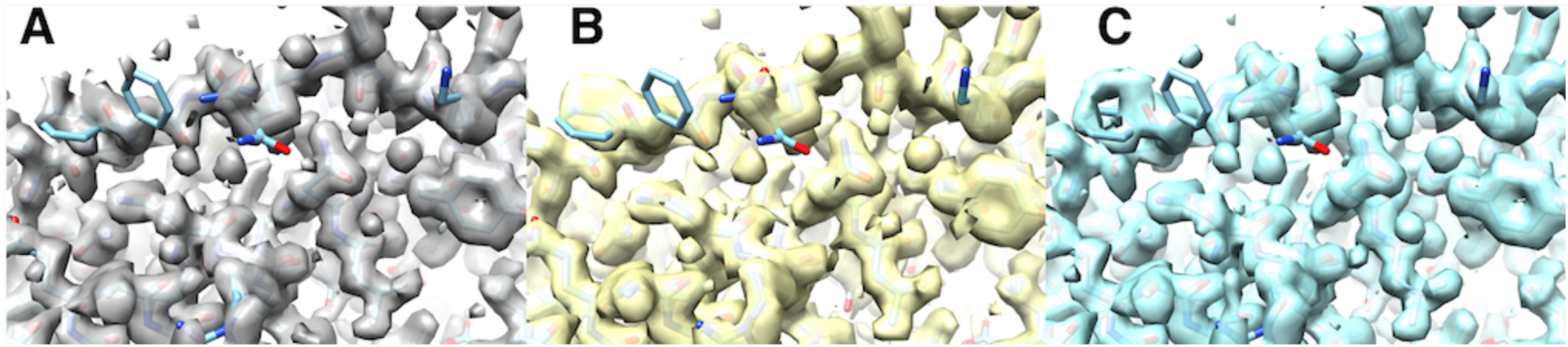
Histogram matching of emd-20026. A. Average of half-maps, sharpened to match density-modified map in C. B, histogram-matched version of map in A, sharpened as in A. C, Density-modified map. All contours set to enclose equal volumes. Maps masked around atoms in model.

**Supplementary Fig. S3.**
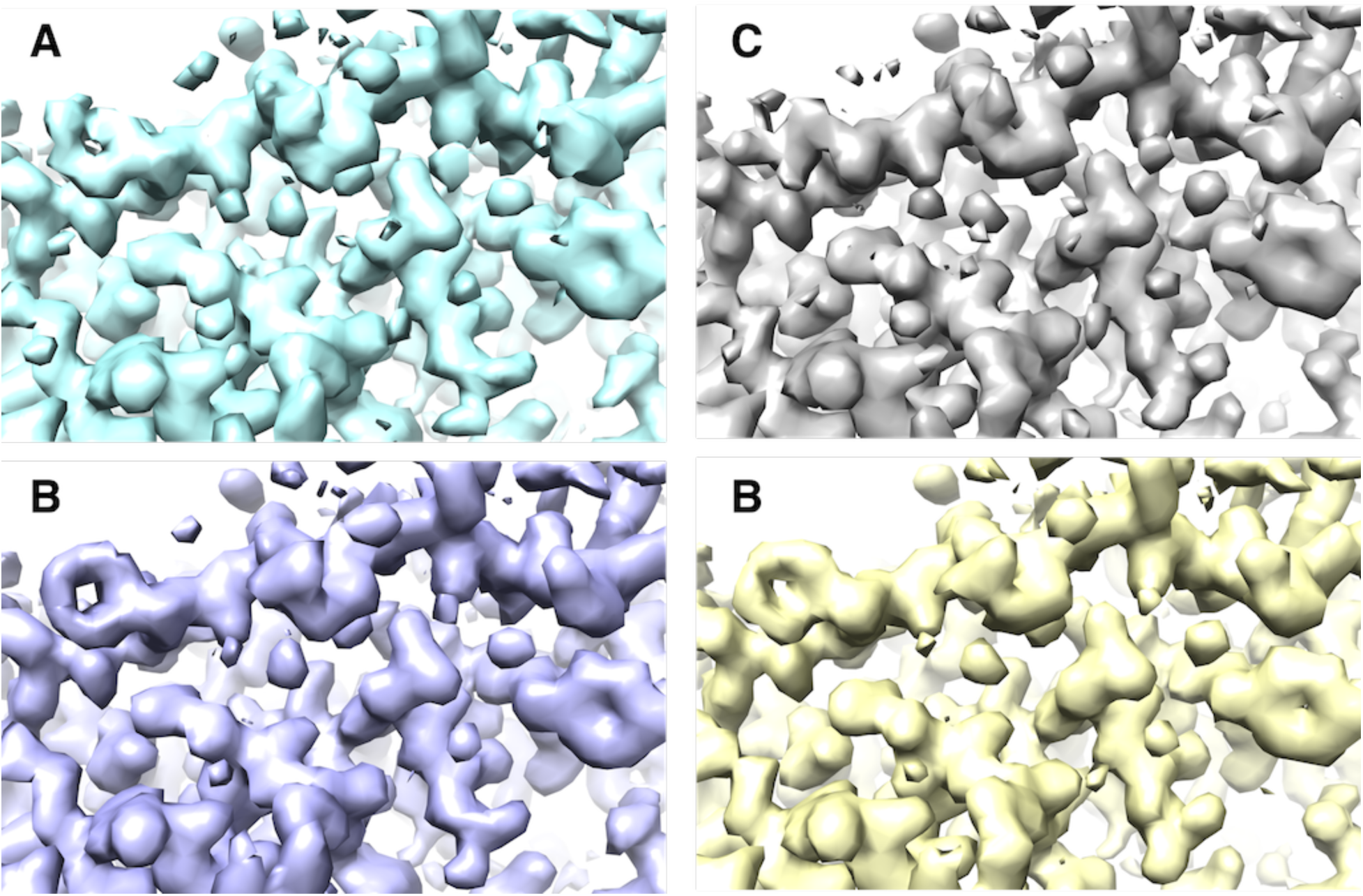
Comparison of half-maps 1 and 2 for density modification shown in Fig. S2. All maps masked and sharpened to match density-modified map in Fig. S2C. A and C, half maps 1 and 2. B and D, density-modified half-maps 1 and 2.

**Supplementary Fig. S4.**
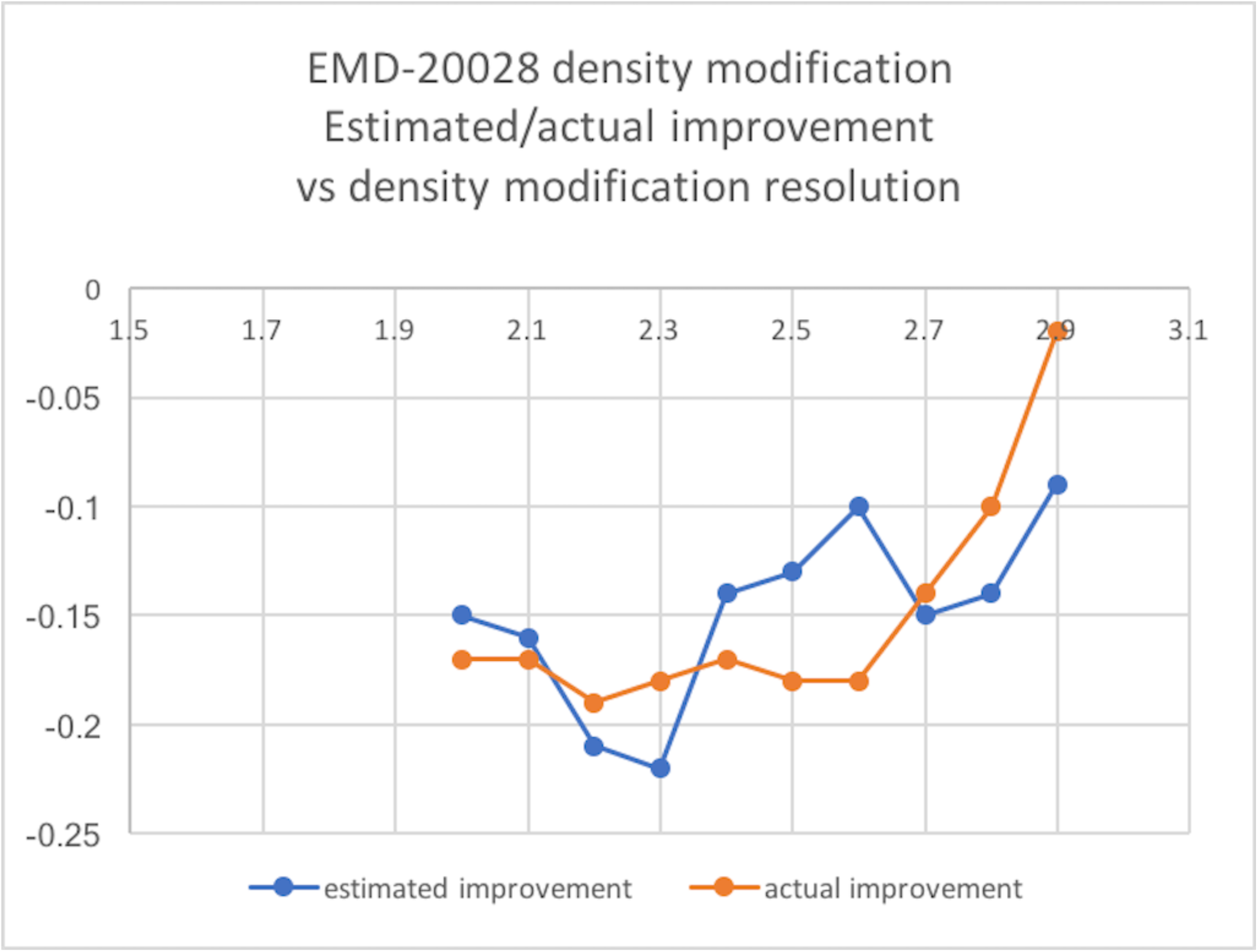
Effect of density modification resolution on actual and estimated change in resolution where FSC is ½. Half-dataset maps for EMD-20028 were density modified as in Fig. 1, but varying the resolution for density modification from 2 to 2.9 Å. The estimate change in resolution (calculated from the error analysis) and the actual change in resolution (calculated from the FSC to the high-resolution EMD-20026 map) are shown.

**Supplementary Fig. S5.**
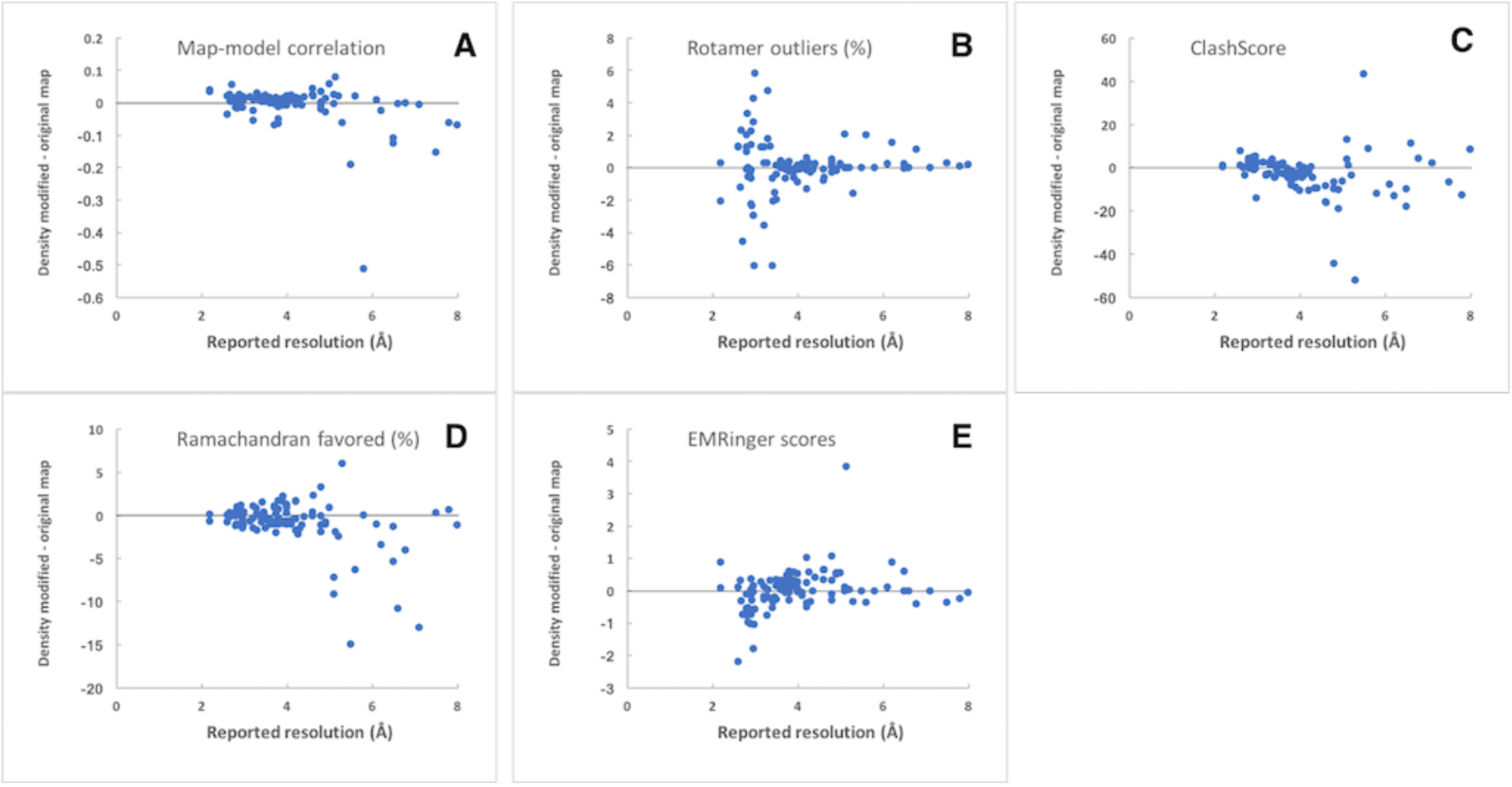
Change in model/map metrics after density modification. Analyses are plotted as a function of resolution for the 104 datasets shown in Fig. 2. A. Map-model correlation, B. Rotamer outliers, C. ClashScore, D. Ramachandran percentage in favored region, E, EMRinger scores.

**Supplementary Fig. 6.**
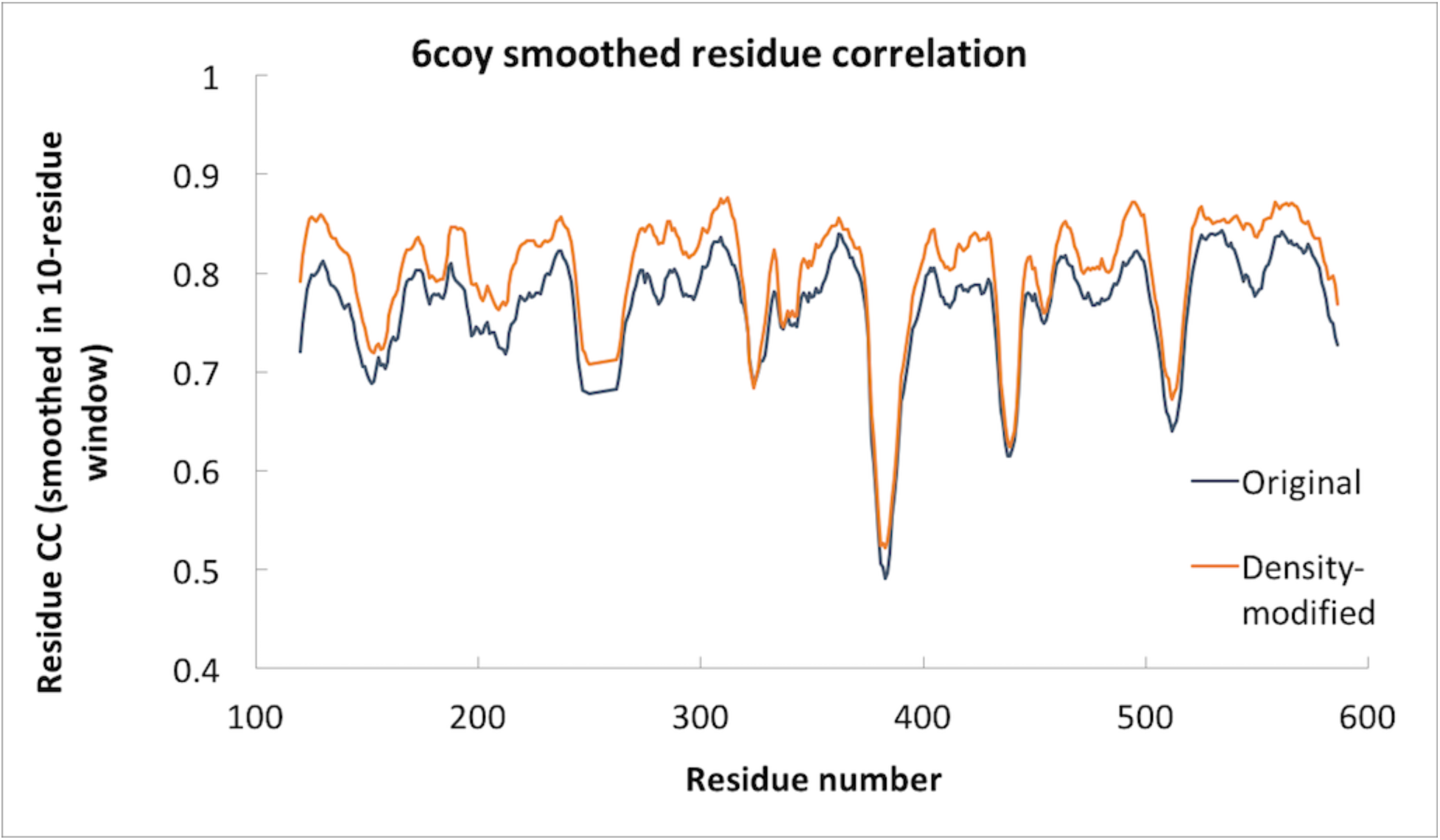
Map-model residue correlation smoothed in 10-residue windows for EMD-7544 (PDB entry 6coy).

**Supplementary Fig. 7.**
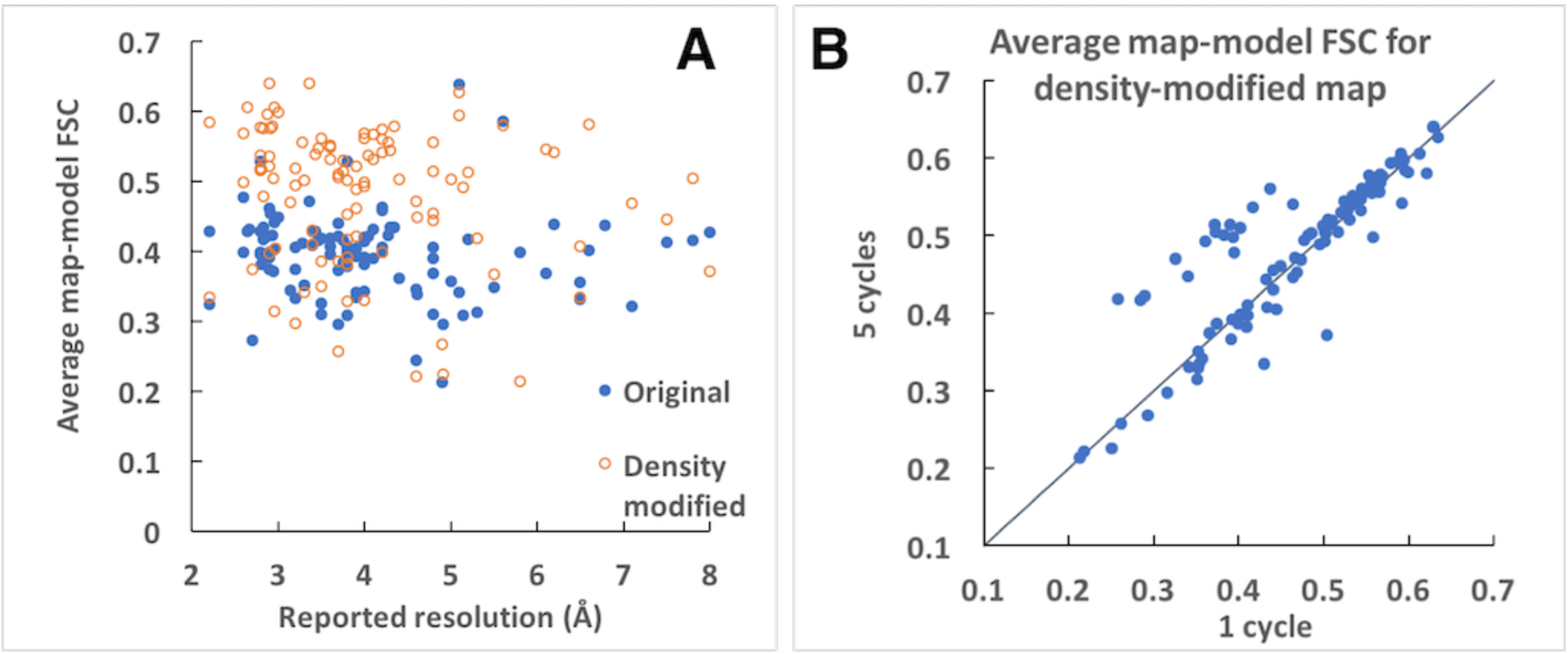
Effect of applying 5 cycles of density modification vs 1. A. Average map-model FSC after 5 cycles of density modification (compare with Fig. 2B). B. Change in average map-model FSC between one cycle of density modification and 5 cycles.

**Supplementary Figure S8.**
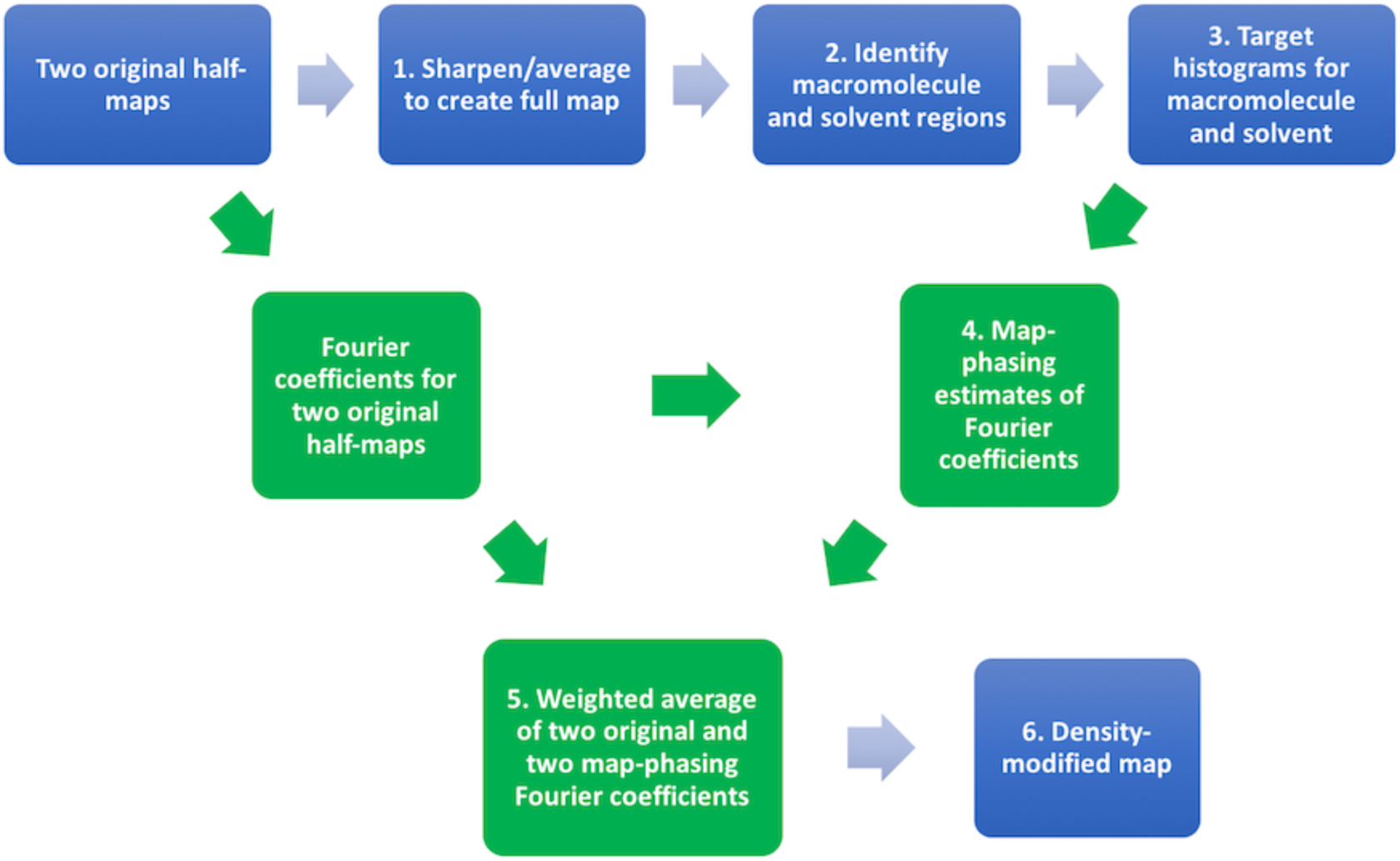
Flow chart of density modification process (see text). Operations in real-space are in blue and those in reciprocal-space are in green.

